# Distraction Impact of Concurrent Conversation on Event-Related Potential Based Brain-Computer Interfaces

**DOI:** 10.1101/2024.04.29.591793

**Authors:** Minju Kim, Sung-Phil Kim

**Affiliations:** Department of Biomedical Engineering, Ulsan National Institute of Science and Technology, Ulsan, Republic of Korea

**Keywords:** electroencephalogram, brain-computer interface, event-related potentials, visual attention, cognitive distractions

## Abstract

**Objective:** This study investigates the impact of conversation on the performance of visual event-related potential (ERP)-based brain-computer interfaces (BCIs), considering distractions in real life environment. The research aims to understand how cognitive distractions from speaking and listening activities affect ERP-BCI performance.

**Approach:** The experiment employs a dual-task paradigm where participants control a smart light using visual ERP-BCIs while simultaneously conducting speaking or listening tasks.

**Main results:** The findings reveal that speaking notably degrades BCI accuracy and the amplitude of ERP components, while increases the latency variability of ERP components and occipital alpha power. In contrast, listening and simple syllable repetition tasks have a lesser impact on these variables. The results suggest that speaking activity significantly distracts visual attentional processes critical for BCI operation

**Significance:** This study highlights the need to take distractions by daily conversation into account of the design and implementation of ERP-BCIs.

## 1. Introduction

One of the frequently utilized brain signals for non-invasive brain-computer interfaces (BCIs), which serve as a communication channel with external devices solely based on brain signals [1–3], is event-related potentials (ERPs) extracted from the electroencephalogram (EEG). ERP-based BCIs (denoted as ERP-BCI hereafter) commonly employ a task paradigm known as the oddball paradigm, where infrequent target stimuli and frequent non-target stimuli are presented randomly. The target stimuli induce P300, also denoted as P3, one of the ERP components that ERP-BCIs mainly harness [4–6].

P3 has been suggested to reflect attentional processes [7– 9]. Larger P3 amplitudes are elicited by attended stimuli than unattended ones [10,11]. In contrast, P3 amplitudes decrease when memory or perceptual processing loads increases [12]. In dual-task scenarios where an oddball task is performed as a secondary task along with an independent primary task, P3 amplitudes vary according to the cognitive demands required for the primary task: the more demanding the primary task is, the more decreased the amplitude is [13–16]. P3 consists of subcomponents: the P3a subcomponent, elicited by novel stimuli, decreases in amplitude with reduced attention due to habituation, while P3b, associated with event categorization, decreases in amplitude with less focus on categorization tasks [8].

Another ERP component prevalently adopted by ERP-BCIs is N200, or N2 [17–21]. Within the context of ERP-BCIs, N2 is primarily distributed over frontocentral or parieto-occipital regions, particularly observed in the occipital region for visual ERP-BCIs. This posterior N2 is affected by the attentional state, similarly to P3 [22,23]. Therefore, various factors related to attentional processes, such as cognitive loads and distraction, can influence the performance of ERP-BCIs. This characteristic of ERP-BCIs should be addressed when deploying ERP-BCIs to everyday life, where people encounter numerous distractions that would vary cognitive loads dynamically.

By modulating mental workload using tasks designed for cognitive research, such as the N-back, mental arithmetic, and dichotic listening tasks, studies have demonstrated that assignment of mental workload to these primary tasks degrades the performance of ERP-BCIs [24–27]. While these tasks can offer well-controlled experimental designs for the impact of mental workload on ERP-BCIs, they are rather far from everyday activities. To investigate the effect of cognitive loads and distraction on ERP-BCIs in daily lives, we should consider distraction factors encountered in everyday activities. One of the everyday activities likely to influence the use of ERP-BCIs by prompting diverse cognitive processes is a daily conversation, encompassing speaking and listening. Due to a limited capacity of human information processing, the cognitive demands of speaking and listening may interfere with the performance of ERP-BCIs. Ergonomic studies on the distracting effect of conversation over the phone on driving or other cognitive tasks revealed that talking on the phone, both handheld and hands-free, diminishes driving performance [28,29]. Moreover, the distraction effect during driving was more dependent on the demands of conversation than on whether using a hands-free phone or not [30].

However, to the best of our knowledge, the effect of conversation on ERP-BCIs has not been explored. Several studies on the influence of listening to speech or music on the ERP-BCI performance have been undertaken, but with inconsistent findings. For instance, engaging in an auditory task of counting the occurrence of a target word increased mental workload leading to the decreased BCI performance [31], whereas attending to background speech did not revealed significant effects on ERP-BCI performance [32]. On the other hand, the effect of speaking on ERP-BCI has yet to be reported. Only a study unrelated to BCIs has shown decreases in P3 amplitudes to visual targets when a concomitant speaking task was conducted [33]. Therefore, further research is needed to understand the effect of speaking and listening on ERP-BCI more clearly.

Visual attention plays a crucial role in ERP-BCI as using visual stimuli outperforms using other sensory stimuli for ERP-BCIs [34–38]. As such, evaluating the degree of visual attention allocated to ERP-BCIs would allow us to understand how conversation tasks impact BCIs. Alpha oscillations observed in the occipitoparietal region have been associated with visual attention [39–42]. Studies have indicated that the enhanced alpha oscillations reflect the inhibition of visual processing, while the suppression of alpha oscillations reflects a state of heightened visual attentional engagement and active visual processing [39,43–45]. Therefore, monitoring alpha oscillations may provide valuable insights into investigating the distraction effect of speaking and listening on ERP-BCI.

In this study, we aimed to investigate the effect of speaking and listening in the context of daily chats on ERP-BCI. We designed a dual task where participants controlled a smart light using ERP-BCI while speaking or listening during conversations at the same time. In addition to the speaking and listening task, a syllable repetition task was also designed to compare the effect of simple vocalization with that of conversational speaking. We examined the BCI performance, ERP components and occipital alpha power in order to investigate the effect of each dual task. We hypothesized that speaking and listening would degrade the BCI performance, decrease the amplitude of ERP components and increase occipital alpha power, but syllable repetition would not.

## 2. Materials and Methods

### 2.1 Participants

Thirty-six healthy undergraduate students (16 females, mean age 19.86 ± 2.02 years old) were recruited for the study. All participants had normal or corrected-to-normal vision, and none of them had a history of neurological disorders. They gave informed written consent before the experiments. This study was approved by the Institutional Review Board of UNIST (UNISTIRB-21-22-A) following the declaration of Helsinki.

### 2.2 Stimuli

The design of visual stimuli for ERP-BCIs was adopted from previous studies [46,47] (Figure 1(A)). As visual stimuli, four blue squares each containing an icon indicating each of the four functions of the smart light to be controlled were presented on the display’s four corners. The four functions included light on, light off, lowering brightness and color change. In each trial, one of the squares flashed by changing its color to light green. A square flashed for 67 ms, with an inter-stimulus interval (ISI) of 67 ms. Each square flashed at a time in a random order within a sequence of four trials. Ten such sequences were presented consecutively, forming of a single block. Thus, the presentation of each square was repeated ten times for a total of forty trials per block. In the beginning of a block, one of the four squares was indicated as a target stimulus by a red border. Participants were instructed to focus on the target stimulus and count the number of times it flashed.

**Figure 1.**
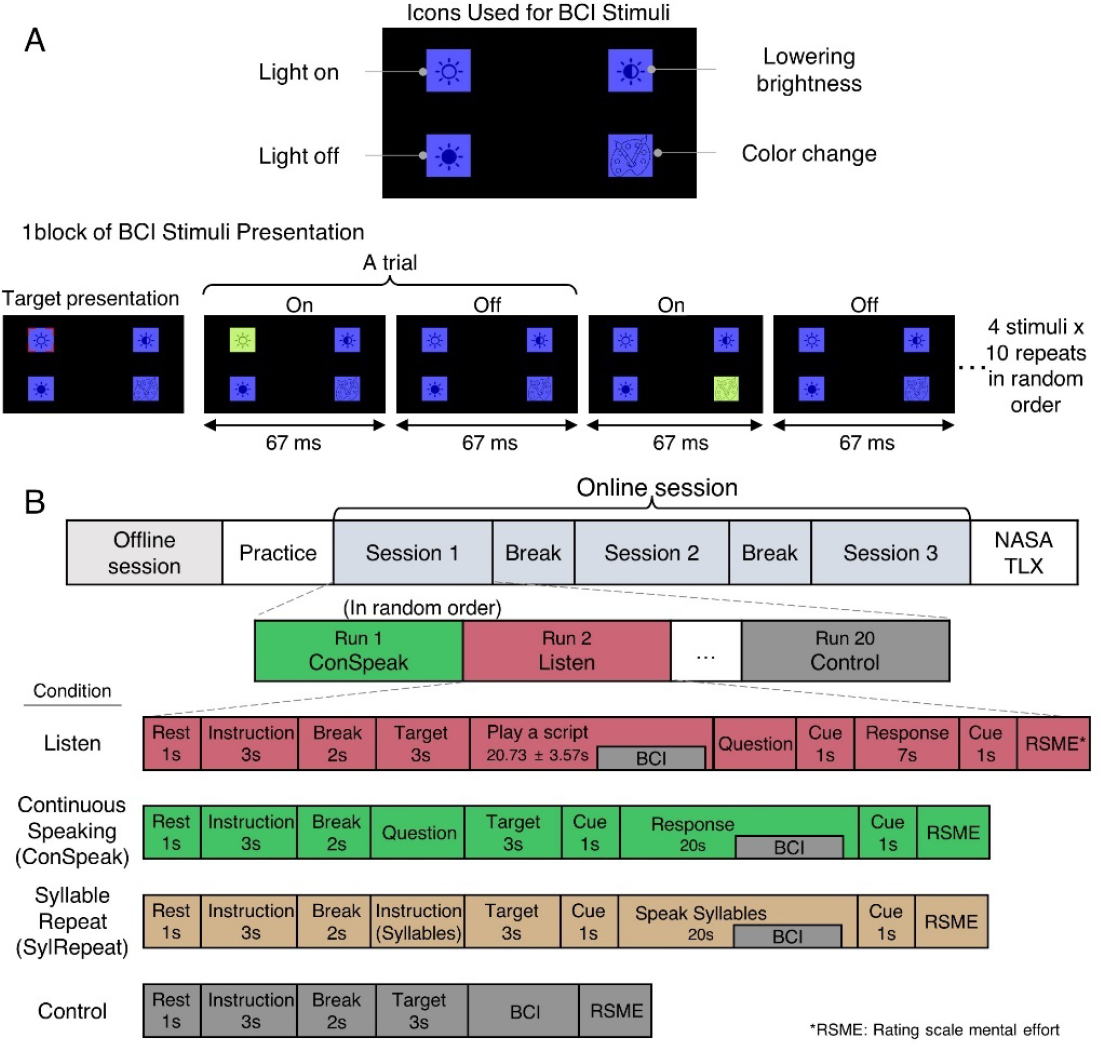
Visual stimuli for ERP-BCIs and the experimental protocol. (A) Visual stimuli design and one block of ERP-BCI stimuli presentation. Once the BCI control task began, four blue squares were presented and a target was marked by a red L-shaped symbol on each corners of the icon. Each stimulus indicated the function of a smart light: light on, light off, lower brightness, and color change. A stimulus was presented by changing its color into light green for 67ms, followed by an inter-stimulus interval of 67ms. Each stimulus was randomly repeated 10 times, resulting in a total of 40 trials. (B) The experiment consisted of offline and online sessions. The offline session comprised 30 blocks of BCI stimuli presentation without feedback. In the online session, participants controlled ERP-BCIs to operate the smart light under four conditions given in a random order: Listen, Continuous speaking (ConSpeak), Syllable repeat (SylRepeat), and Control. Every run in the session started with 1s of resting and an instruction informing the task to be performed. Participants performed either dual tasks (Listen, ConSpeak and SylRepeat) or a single task (Control) (See Section 2.3.3 for more information about each task). The run ended with rating scale mental effort (RSME) survey. Totally three sub-session constituted the online session and each sub-session was composed of 20 runs. After the whole experiment was completed, participants responded to NASA TLX questionnaire.

### 2.3 Experimental procedure

The experiment consisted of offline and online sessions (Figure 1(B)). The offline session was designed for collecting data to train a classification model. Then, using the trained model, participants controlled the smart light through ERP-BCIs while performing a dual task at the same time in the online session. Each participant completed the offline session first and conducted four blocks of practice before the online session. Details of each session are described in the following sections 2.3.1. and 2.3.2.

#### 2.3.1 Offline session

The offline session was composed of thirty blocks of stimulus presentation. No feedback about target selection was provided after each block. There was a short break (3s) between blocks. Participants solely performed the oddball task without engagement in any other task.

#### 2.3.2 Online session

The online session consisted of three sub-sessions, each comprising twenty runs. Each run underwent in one of the four conditions: Listening (Listen), Continuous Speaking (ConSpeak), Syllable Repeat (SylRepeat), and Control. The description of each condition will be provided below. Participants controlled ERP-BCIs under each condition five times in a random order per sub-session. As such, there were fifteen runs total per condition in the online session. A short break was provided between sub-sessions where participants voluntarily started the next session anytime by pressing a space bar.

In the online session, a single run was started with instructions informing participants which of the four tasks associated with the four conditions above they should perform. Also, participants were informed that the smart light beside them would be controlled as a result of using ERP-BCIs. The smart light was positioned at the left side of participants, situated between them and the monitor, so they were able to see the operation of the light. After each block of stimulus presentation, the real-time feedback was directly provided to participants by operating one of the four functions of the smart light according to decoding output from ERP-BCIs. At the end of a run, participants responded to the Rating Scale Mental Effort (RSME) [48] on screen, where a line with a scale ranging from 0 to 150 was displayed, alongside nine anchor points with verbal descriptors, including “absolutely no effort”, “almost no effort”, ”a little effort”, “some effort”, “rather much effort”, “considerable effort”, “great effort”, “very great effort”, and “extreme effort”. Participants marked a point by clicking on the scale on the display using a mouse to report how much mental effort was required to perform the task. After all sub-sessions were finished, participants completed the NASA TLX questionnaire [49].

#### 2.3.3 Task

Participants performed dual tasks under the three conditions of Listen, ConSpeak, and SylRepeat, including a primary task specifically designed for the corresponding condition and a secondary BCI control task with the oddball paradigm. In contrast, they performed only the BCI control task under the Control condition. The initiation timing of the BCI control task in the course of the primary task was pseudo-randomized but adjusted not to start too early or too late (Figure 1(B)). This adjustment was for obtaining the consistency in the dual task environment, where two tasks had to be performed simultaneously, ensuring that participants were not engaged exclusively in the BCI task at any point under dual task conditions. The tasks in each condition are described below.

##### 1) Listen

An audio script was played through two audio speakers that were positioned beside the monitor. Scripts used in the experiments were edited based on passages from Korean high school textbooks. Fifteen different scripts were prepared with the mean length of 20.73 ± 3.57 s. In the middle of playing a script, the BCI control task started at a random moment. Participants had not known a priori when the BCI control task would start, but the actual start timing was maintained identical for all participants, in order to ensure that all participants were engaged in the BCI task while paying attention to the information relevant to questions to be asked after the task. Following the script, a question about the contents of the script was given, and participants answered the question verbally. The questions were prepared to include words mentioned in the script as a correct answer so that participants needed to avoid missing contents as much as possible.

##### 2) ConSpeak

Following the instruction about the task, a question asking the opinion of participants was given. Questions followed an agree/disagree style such as, “Can government policies increase the fertility rate?” or “Is it okay to tell a white lie?”. The BCI control task started after the question was given. Participants were required to give their opinion verbally for 20 s until an auditory cue was given. They were asked to speak as constantly as possible without a long pause.

##### 3) Syllable Repeat (SylRepeat)

After the instruction of the task was given, another instruction informing which kind of syllables to speak repeatedly was additionally provided. Syllables to vocalize were designed using three simple Korean syllables: “Ga Na Da,” “Ma Ba Sa,” or “Ta Pa Ha.” These syllables were designed to be meaningless and as simple as possible so that little effort was required to pronounce them. Participants were given one of these syllable sets randomly and pronounced the syllables loud repeatedly until an auditory cue was given for 20 s.

##### 4) Control

Without any other concurrent tasks, participants were instructed to perform the BCI control task. It took 11.3s, including stimulus presentation and feedback presentation, to complete the BCI control task, which was consistent over all conditions.

### 2.4 EEG recordings

EEG was acquired from 31 Ag/AgCl scalp active wet electrodes (FP1, FPz, FP2, F7, F3, Fz, F4, F8, FC5, FC1, FCz, FC2, FC6, T7, C3, Cz, C4, T8, CP5, CP1, CPz, CP2, CP6, P7, P3, Pz, P4, P8, O1, Oz, and O2) according to the International 10/20 system using a commercially available amplifier (actiCHamp plus, Brain Products GmbH, Germany). The ground and reference were attached to the left and right mastoids respectively. The sampling rate was 500 Hz, and electrode impedance was kept below 10 k Ω . Additional passive electrodes were attached above the upper lip and below the lower lip to capture muscle activity from the left orbicularis oris [50].

### 2.5 EEG processing for ERP-BCI

#### 2.5.1 EEG Preprocessing

EEG signals were high-pass filtered (0.5 Hz ∼) first and low-pass filtered (∼ 50 Hz) afterward using finite impulse response (FIR) filters. Bad channels were identified and removed based on correlations between channels. When over 70 % of other channels were weakly correlated (<0.4), that channel was deemed to be a bad channel [46]. Common average reference (CAR) was applied for re-referencing, and artifacts were rejected by artifact subspace reconstruction (ASR) [51]. The continuous EEG signal was segmented to epochs from 200 ms before to 600 ms after stimulus onset and baseline-corrected by subtracting the mean of baseline (-200 ms ∼ 0 ms). Epoched data was downsampled to 100 Hz before feature extraction.

#### 2.5.2 Feature extraction

We applied the discriminative spatial pattern (DSP) method for feature extraction [52]. DSP, inspired by the Fisher discriminant analysis, was introduced as a spatial filter for the extraction of a feature space to maximize differences between classes (i.e., target vs. non-target in our case). Let *X*_*j*_ (*i*) denote the EEG signal in the ith trial of class *j*, comprising an *N* x *T* matrix, where *N* is the number of channels and *T* is the number of samples in an epoch. Let *M*_*j*_ ∈ ℝ^*NxT*^denote the average of *X*_*j*_ (*i*) over trials from class *j*. Let Sw be the within-class scatter matrix and SB be the between-class scatter matrix, given by:

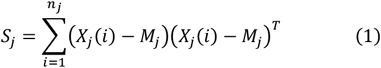

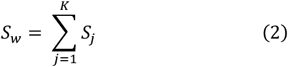

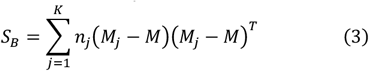

where *M* is the average of all trials from all classes, *n*_*j*_ is the number of trials from class *j* and *K* is the number of classes. Fisher’s linear discriminant criterion is given as:

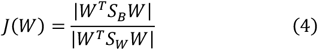

where a matrix *W* projects the data onto low-dimensional spaces to maximize between-class variance and minimize within-class variance. We can find the solution of *W* to maximize *J(W)* by solving the generalized eigenvalue problem,

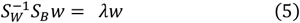

where λ is the eigenvalue and *w* is the eigenvector. The eigenvectors *w*’s constitute the columns of *W* and plays a role of spatial filtering. We also regularized *S*_*w*_ as,

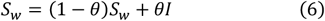

with *θ* = 0.1 [52] where *I* indicates an identity matrix. Using the projection matrix *W*, a single-trial (*i*) is transformed into,

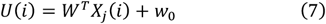

*U* is the transformed signal of *N*_*f*_ x *T* where *N*_*f*_ is the number of spatial filters (i.e., the number of eigenvectors used). *w*_0_ is a bias term calculated as -*W*^*T*^*M* . In this study, we determined the number of spatial filters for each participant from the 5-fold cross-validation within the training data. The number of spatial filters that led to the highest cross-validation accuracy was used for feature extraction. Consequently, the number of spatial filters used in the online session was 7.97 ± 3.17 on average. The transformed signals (*U*(*i*)) within a window of 0 ms to 400 ms were used as features after downsampling with a factor of 5, resulting in the feature dimension of 40 × *N*_*f*_ . Note that we opted for DSP over well-known the common spatial pattern (CSP) method widely used for BCIs, because DSP was designed to separate non-oscillatory event-related potentials by maximizing amplitude differences while CSP aimed at maximizing differences in variance between signals.

#### 2.5.3. Classification

The support vector machine (SVM) [53] with a sigmoid kernel was used to classify target and non-target ERPs. The SVM classifier was learned with the training data, which comprised 300 samples of the target and 900 samples of non-target class (from 10 (30) trials / block x 30 blocks for target (non-target)). In the online session, a target stimulus in a given block was classified among four stimuli as follows:

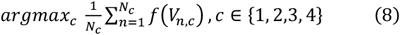

where *V*_*n*,*c*_ denotes a feature vector in the *n*-th trial of the *c*-th stimulus, and *f* indicates an SVM classifier producing SVM decision values that represent distances from the decision boundary to *V*_*n*,*c*_ . A more positive decision value can be regarded as a higher chance of belonging to the target class. *N*_*c*_ is the number of trials of the presentation of the *c*-th stimulus per block (i.e., *N*_*c*_ = 10). The accuracy of ERP-BCI was measured by a ratio of the number of blocks producing correct command to that of the total number of blocks.

### 2.6 Data analysis

In addition to the evaluation of the online BCI performance, we conducted post-hoc analyses of ERP, alpha oscillations and SVM decision values using the test data from the online session among all four experimental conditions (Listen, ConSpeak, SylRepeat, and Control).

#### 2.6.1 ERP

We analyzed ERP peak and latency to examine the effect of conditions on ERP components. We also examined ERP latency variability as an indicator of distraction that could result in temporal variations in cognitive processing in the oddball task. The quality of ERP data was assessed whether potential artifacts induced by dual tasks (e.g., mouth movements) affected ERP data.

From averaged ERPs across the whole trials in each condition for each participant, peak amplitudes and latencies of P3a, P3b, and occipital N2 components were estimated. The electrodes of interest in our study included Fz, FC1, FC2, and Cz for P3a, P3, Pz, and P4 for P3b, and O1, Oz and O2 for occipital N2. According to the previous studies, we set time windows specific to each component: 100∼300 ms for P3a and N2, and 250∼450 ms for P3b [17,54]. In these windows, we identified the peak latency as the timing of the maximum (P3a and P3b) or minimum (N2) value. Once we identified the peak latency of each ERP component for each participant, we estimated a peak amplitude as the mean amplitude averaged from 25 ms before to 25 ms after the peak latency. Note that we employed the mean amplitude rather than a single peak amplitude because the mean amplitude is more robust to noise than peak amplitude. Then, we examined the effect of the experimental condition on the peak amplitude for each ERP component. In the following analyses, we used the absolute value of the peak amplitude to facilitate a comparison of how the peak amplitude of each component changed.

Differences in ERP amplitudes among the conditions can arise not only from changes in the amplitude itself but also from latency variability across single trials. Therefore, we also examined the latency variability of ERP components. For each ERP component, we first calculated the cross-correlation with varied time lags between a template ERP and a single-trial ERP in all 150 target trials of the online session in each condition. The template ERP was separately determined by averaging EEG data in 300 target trials from the offline session. The cross-correlation curves as a function of time lag obtained from each electrode were averaged across electrodes of interest of respective ERP components. A latency shift per trial was calculated at which the electrode-averaged cross-correlation curve was maximized [55]. The latency variability was defined as the standard deviation of the latency shifts across all 150 target trials. The latency shifts exceeding ± 50 ms of time lag were not included for standard deviation estimation. We compared the latency variability among the conditions for each ERP component.

Due to potential imprecision in ERP component measurements caused by artifacts from facial movements in ConSpeak and SylRepeat, the standardized measurement error (SME) was applied to assess the quality of ERPs. SME was developed to assess the quality of ERP measurements, such as peak amplitude and peak latency [56]. SME is defined as the standard error of ERP measurements, where ERPs are measured by averaging epoched EEG data, indicating the consistency of ERP measurements across observed EEG data. In this study, the bootstrapping method was employed to estimate the standard error by repeatedly generating ERP measurements from the entire data, to obtain a sampling distribution of ERP measurements [56]. For each condition, 150 trials were randomly sampled with replacement across which EEG data were averaged to obtain an ERP measurement. The peak amplitude and latency were estimated from the ERP measurement following the aforementioned process. This sampling and estimating process was repeated 1,000 times. The standard deviation of resulted peak amplitude and latency was calculated as the SME, respectively. The SME values for each ERP component were computed at each corresponding electrode of interest.

#### 2.6.2 Alpha oscillations

We calculated the power of alpha oscillations to assess variations of alpha oscillations across the conditions. Alpha band power was calculated from a time segment of 5.4 s in which the BCI control task was performed for each condition.

Power spectral density (PSD) was estimated from EEG signals using Welch’s method with a Hanning window (500 ms) and 50 % overlap. In order to calculate alpha power, an area of the PSD within the frequency range of 8 Hz to 12 Hz was integrated. We defined alpha modulation in the dual tasks as the ratio of alpha power in each block from dual tasks to that of the Control condition, averaged across all blocks:

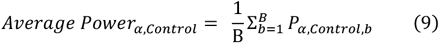

where *B* is the total number of blocks (i.e., *B* = 15) and *P*_*α, Control*,*b*_ is the alpha power in the *b*-th block under the Control condition. Then, the alpha power modulation level was defined as:

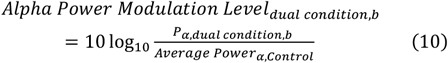

where *P*_*α*,*dual Condition*,*b*_ is the alpha power in the *b*-th block under a specific dual task condition. The alpha power modulation level was estimated for every block under dual conditions at the occipital channels (O1, Oz, O2) and averaged across those channels. The average of alpha modulation level across blocks in dual task conditions was used for statistical tests.

#### 2.6.3 Analysis on classification outputs

To test the impact of dual tasks on data separability between target and non-target trials, we conducted a separability analysis on SVM classification outputs. We used *d’* (d prime) as a separability metric to quantify the degree of separation between the SVM decision values, the distance from the decision boundary to data points, of target and non-target trials. *d’* was calculated as follows:

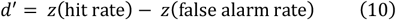

where *z*(*⋅*) represents z-transformation. Hit rate is the proportion of target trials classified as target, and False alarm rate is the proportion of nontarget trials classified as target. *d*’ values were compared across conditions to assess how cognitive distractions influenced the separability. This separability analysis allowed us to evaluate how distractions reflected in the ERP features influenced the SVM’s ability to discriminate between target and non-target ERP features.

#### 2.6.4 Statistical tests

We used Friedman test as the dependent variables, including BCI accuracy, subjective assessments, ERP amplitude, ERP latency variability and alpha modulation level, did not follow a normal distribution (Kolmogorov-Smirnov Test, p < 0.05). For the post-hoc test, Tukey’s honestly significant difference (HSD) was used. The multiple comparison for of ERP amplitudes for different electrodes was corrected using the false discovery rate (FDR) with α < 0.05 [57]. We applied the Spearman correlation coefficient when measuring correlations between ERP amplitudes and the accuracy, between latency variability and the accuracy, and between the alpha modulation level and the accuracy, respectively.

## 3. Results

### 3.1 BCI performance

The online accuracy of ERP-BCIs under the four conditions, Listening, ConSpeak, SylRepeat and Control, was evaluated (Figure 2(A)). The mean online accuracy was the highest in Control condition (96.82 ± 4.66 %, MEAN±STD), followed by SylRepeat (95.90 ± 6.47 %) and Listening (93.05 ± 9.36 %), and the lowest in ConSpeak condition (88.11 ± 14.11 %). Friedman test showed a significant effect of condition on accuracy (*χ*^2^(3) = 17.54, *p* < 0.0001). The post-hoc test revealed that the mean accuracy in ConSpeak condition was significantly lower than those of Control (*p* = 0.0007) and SylRepeat conditions (p = 0.0042). The difference in the mean accuracy between ConSpeak and Listening was marginally significant (*p* = 0.062). There was no difference among other condition pairs (*ps* > 0.1).

**Figure 2.**
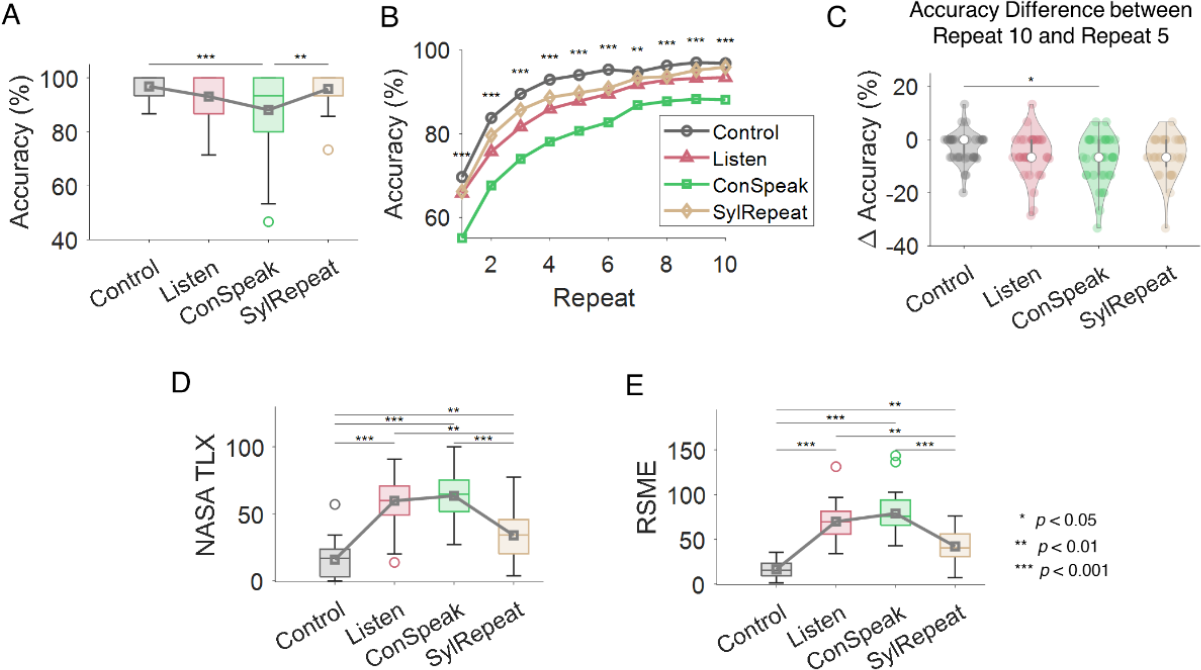
BCI performance and mental effort assessment. (A) Online BCI accuracy in each condition across all 36 participants. The empty circles indicate outliers, and the gray square shows the average accuracy. The boxes denote 25% and 75% percentiles. (B) BCI accuracy in each condition as a function of the number of repetitions. The asterisks represent the significance of the difference in accuracy between the conditions at each repetition (Friedman Test). (C) The accuracy difference from 10 repetitions to 5 repetitions. Single colored-dots denote participants and empty dots show median values. (D) The subjective level of workload (NASA TLX score) in each condition. This score was measured after all experimental sessions were completed. (E) The subjective level of mental effort (Rating scale mental effort, RSME) at each condition. This scale was measured at the end of every run. Statistical significance was evaluated by Friedman test with Tukey’s HSD post-hoc test, *** *p* < 0.001, ** *p* < 0.01, * *p* < 0.05.

Next, we performed an offline analysis to examine the effect of condition on accuracy according to the number of stimulus repetitions (Figure 2(B)). We found significant effects of condition for all repetitions from one to ten (Friedman test, *p* < 0.005, FDR-corrected). Further, we examined if BCI performance degradation due to fewer repetitions differed among conditions. To that end, we calculated a difference of accuracy between ten and five repetitions and compared it among conditions (Figure 2(C)). Friedman test revealed that there was a significant effect of condition (*χ*^2^(3) = 9.05, p = 0.0287) on the performance difference. The post-hoc test showed that the performance degraded significantly more in ConSpeak than in Control (*p* = 0.0178), while no significant differences between other condition pairs were observed (*p* > 0.1). It showed that the distraction effect by continuous speaking sustained regardless of the number of repetitions and increased when the number of repetitions to obtain ERPs was relatively small.

### 3.2 Subjective assessment

The mental workload and effort required for participants to perform the task were assessed using NASA TLX (Figure 2(D)) and RSME (Figure 2(E)). NASA TLX was measured per condition after all experiments were finished, while RSME was measured at the end of every run. RSME values were averaged across runs per condition. The assessment results showed that NASA TLX and RSME were both the highest in the ConSpeak condition, followed by Listen, SylRepeat, and Control in an order. Friedman test showed a significant effect of condition on both NASA TLX (*χ*^2^(3) = 77.29, *p* < 0.0001) and RSME (*χ*^2^(3) = 94.43, *p* < 0.0001). The mean NASA TLX score and RSME of Control were significantly lower than those of Listen (*p* < 0.0001), ConSpeak (*p* < 0.0001), and SylRepeat (*p* < 0.005). The mean NASA TLX score and RSME of SylRepeat were significantly lower than Listen (*p* = 0.001) and ConSpeak (*p* < 0.0005). There was no significant difference between Listen and ConSpeak (*p* = 0.9466 for NASA TLX and *p* = 0.4065 for RMSE).

### 3.3 ERP components

The grand mean ERPs across all participants at the electrodes of interest (see Section 2.6.1) are depicted in Figure 3 for all conditions. Figure 3(A) illustrates the difference waveforms between target and non-target ERPs. Figure 3(B) shows the topographies of target ERPs over time from 100 ms to 600 ms poststimulus. These overall ERP patterns indicate that the amplitudes of target ERPs in the ConSpeak condition were relatively decreased compared to other conditions (Figure 3(B)), potentially leading to smaller differences between target and non-target ERPs (Figure 3(A)).

**Figure 3.**
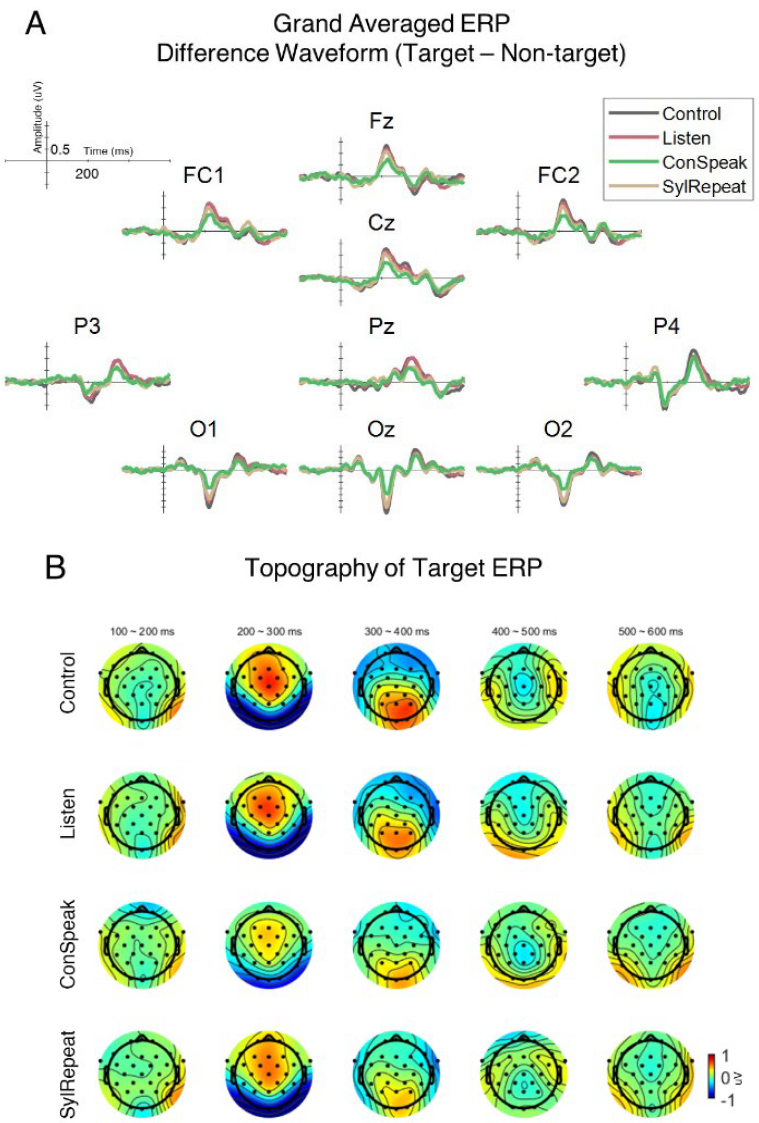
ERP waveform and topography. (A) The difference waveform between grand averaged target and non-target ERPs at Fz, FC1, FC2, Cz, P3, Pz, P4, O1, Oz, O2 under four different conditions. The vertical lines denote the stimulus onset. (B) The topographies of target ERPs over 31 EEG channels under four different conditions. The ERP amplitudes were averaged over the time windows of 100 ms.

#### 3.3.1 Peak amplitude of ERP components

The peak amplitude of ERP components including P3a, P3b, and N2 were compared between conditions at the electrodes of interest for each component (Figure 4(A)). Specifically, we examined P3a at Fz, FC1, FC2 and Cz, P3b at P3, Pz, and P4, and N2 at O1, Oz and O2. Overall, we found significant differences in the peak amplitude of all the ERP components (Friedman test, p < 0.05, Table 1). We also compared the peak latency of ERP components between conditions at the electrodes above, but found no significant difference (Friedman test, *p* > 0.05, Table S1).

**Table 1.** Mean and Friedman test results of ERP peak amplitude

| Channel | Peak Amplitude (uV) | | | | Chi-square | $p$-value (corrected) |
| --- | --- | --- | --- | --- | --- | --- |
|  | Control | Listen | ConSpeak | SylRepeat |  |  |
| Fz | 1.0147 | 0.9111 | 0.6207 | 0.8129 | 17.90 | 0.0005 |
| FC1 | 0.9442 | 0.8900 | 0.5994 | 0.6747 | 20.93 | 0.0001 |
| FC2 | 0.9687 | 0.8340 | 0.5994 | 0.7404 | 19.23 | 0.0003 |
| Cz | 1.0147 | 0.9111 | 0.6207 | 0.8129 | 22.93 | 0.0007 |
| P3 | 0.7464 | 0.7696 | 0.5508 | 0.5508 | 16.90 | 0.0007 |
| Pz | 1.0286 | 0.9488 | 0.6234 | 0.7387 | 26.38 | $1.9824 \times 10^{-5}$ |
| P4 | 1.1076 | 0.8882 | 0.7659 | 0.8420 | 25.83 | $2.0670 \times 10^{-5}$ |
| O1 | -2.1210 | -1.7756 | -1.1645 | -1.7147 | 29.75 | $5.3195 \times 10^{-6}$ |
| Oz | -2.3904 | -1.9813 | -1.4695 | -1.9915 | 29.70 | $5.3195 \times 10^{-6}$ |
| O2 | -2.2736 | -1.8968 | -1.3660 | -1.9834 | 39.79 | $1.1813 \times 10^{-7}$ |

**Figure 4.**
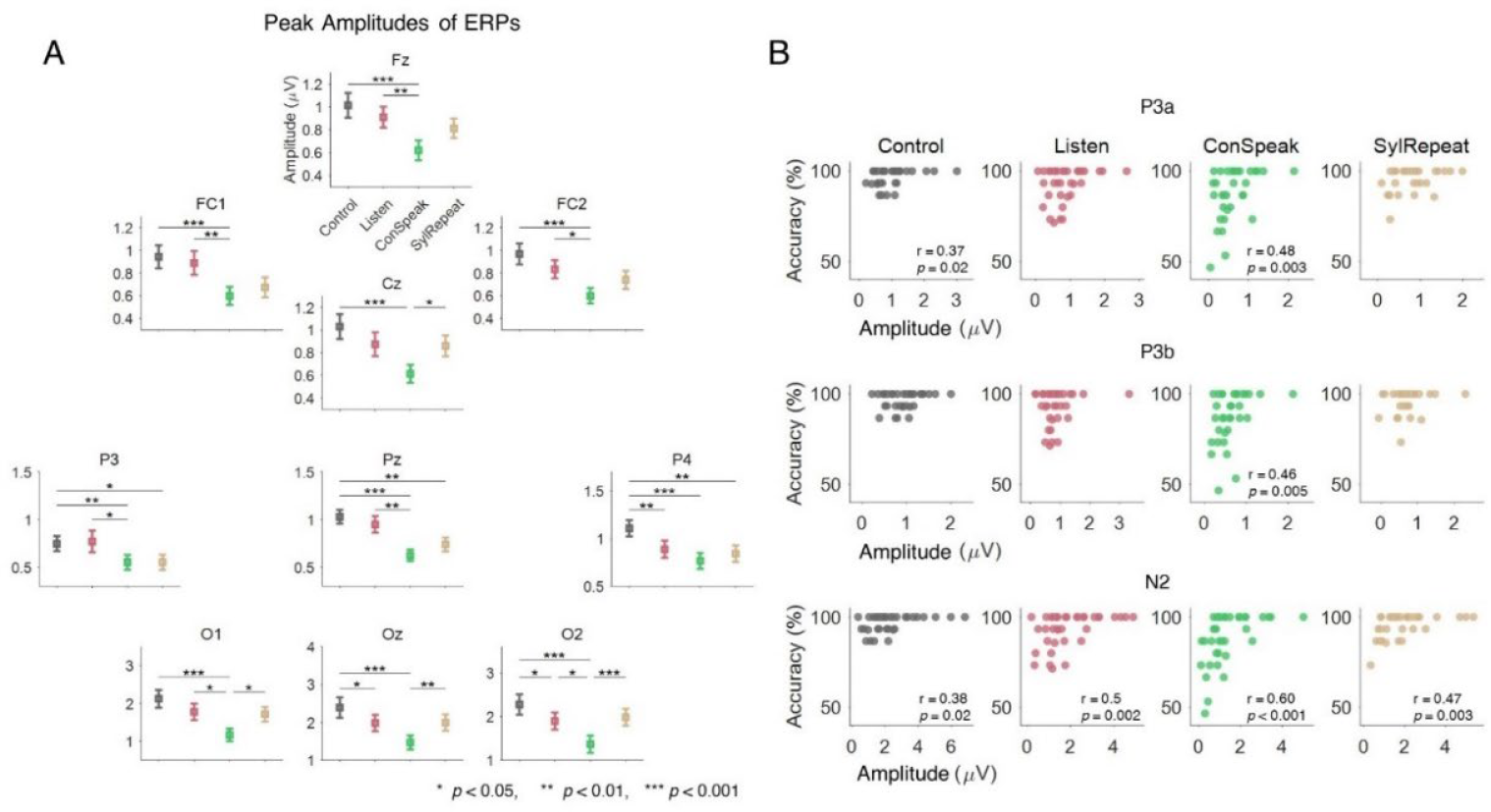
The peak amplitudes of ERP components and their correlations with the accuracy under different conditions. (A) The average absolute peak amplitudes of ERP components at the channels of interest (see the text for details of channel selection). The peak amplitudes were averaged across all 36 participants. The error bars represent the standard error of mean (SEM). The peak amplitudes at Fz, FC1, FC2, Cz were estimated from the time window of 100ms to 300ms, concerning the ERP component of P3a. From P3, Pz and P4, the peak amplitudes were calculated over 250ms to 450ms, concerning P3b. The peak amplitudes at O1, Oz, and O2 were measured from 100ms to 300ms, concerning N2. Statistical significance was evaluated by Friedman test with Tukey’s HSD post-hoc test and corrected using FDR with α < 0.05. *** *p* < 0.001, ** *p* < 0.01, * *p* < 0.05. (B) The relationship between ERP peak amplitudes and BCI accuracies. Single dots represent individual participants. ERP peak amplitudes were averaged across the corresponding channels: Fz, FC1, FC2 and Cz for P3a, P3, Pz, and P4 for P3b, and O1, Oz and O2 for N2. A significant correlation is depicted by the presentation of *p*-value.

In detail, the P3a amplitude in ConSpeak was significantly smaller than those in Control (*p* = 0.0007 for Fz, *p* = 0.0002 for FC1, *p* = 0.0001 for FC2, and *p* < 0.0001 for Cz) and Listen (*p* = 0.0041 for Fz, *p* = 0.0029 for FC1, *p* = 0.0183 for FC2). No significant difference was found between ConSpeak and SylRepeat except for Cz where the amplitude was smaller in ConSpeak than in SylRepeatk (*p* = 0.0183 for Cz).

The P3b amplitude in ConSpeak was significantly smaller than those in Control (*p* = 0.0029 for P3, *p* < 0.0001 for Pz and P4), and SylRepeat (*p* = 0.0138 for P3, *p* = 0.0025 for Pz, and *p* = 0.001 for P4). It was also significantly smaller than that in Listen at both P3 (*p* = 0.0405) and Pz (*p* = 0.0048). At P4, the P3b amplitude in Listen was significantly smaller than in Control (*p* = 0.0021).

The N2 absolute amplitude was significantly smaller in ConSpeak than in SylRepeat *(p* = 0.0188 for O1, *p* = 0.0076 for Oz, and *p* < 0.0001 for O2) and in Control (*p* < 0.0001) at O1, Oz and O2. In addition, it was significantly smaller in ConSpeak than in Listen at O1 (*p* = 0.0188) and O2 (*p* = 0.0161). The N2 amplitude was significantly smaller in Listen than in Control at Oz (*p* = 0.0138) and O2 (*p* = 0.0121).

We also inspected correlations between the peak amplitude and BCI accuracy across individuals (Figure 4(B)). When calculating correlations, we averaged the peak amplitudes over electrodes for respective components (Fz, FC1, FC2, Cz for P3a, P3, Pz, P4 for P3b and O1, Oz, O2 for N2). Significant correlations were observed between the P3a amplitude and accuracy in Control (*r* = 0.3721, *p* = 0.0254) and ConSpeak (*r* = 0.4859, *p* = 0.0027). The P3b amplitude was significantly correlated with accuracy only in ConSpeak (*r* = 0.4619, *p* = 0.0046). The N2 amplitude was significantly correlated with accuracy in all conditions (*r* = 0.3785, *p* = 0.0028 for Control, *r* = 0.5000, *p* = 0.0019 for Listen, *r* = 0.6012, *p* = 0.0001 for ConSpeak, *r* = 0.4760, *p* = 0.0033 for SylRepeat). Thus, the reduction of the peak amplitudes by the dual task influenced individual BCI performance most significantly in the ConSpeak condition.

#### 3.3.2 Latency variability

In examining the latency variability of individual ERP components across different conditions, the following observations were made (Figure 5(A), Table 2). Overall, the latency variability of all three ERP components was the highest in ConSpeak. Specifically, the latency variability in ConSpeak was significantly higher than that in Control for all components, (*p* < 0.0001 for P3a P3b, and N2) or that in Listen for P3a (*p* = 0.0011) and N2 (p = 0.0002). Furthermore, the latency variability in Listen was significantly higher than that in Control for P3b (*p* = 0.0207), and N2 (*p* = 0.0365). Lastly, the latency variability in SylRepeat was higher than that in Control for P3a (*p* = 0.0005) and N2 (*p* = 0.0005), but lower than that in ConSpeak for P3b (*p* = 0.0061) and N2 (*p* = 0.0207).

**Table 2.** Mean and Friedman test results of ERP latency variability

| ERP Component | Latency Variability | | | | Chi-square | $p$-value (corrected) |
| --- | --- | --- | --- | --- | --- | --- |
|  | Control | Listen | ConSpeak | SylRepeat |  |  |
| P3a | 16.3257 | 16.7393 | 17.6236 | 17.1068 | 34.2333 | $2.6532 \times 10^{-7}$ |
| P3b | 15.1855 | 15.7875 | 16.3134 | 15.6014 | 28.9333 | $2.3128 \times 10^{-6}$ |
| N2 | 13.3514 | 14.0599 | 15.4408 | 14.3606 | 48.3000 | $5.5150 \times 10^{-10}$ |

**Figure 5.**
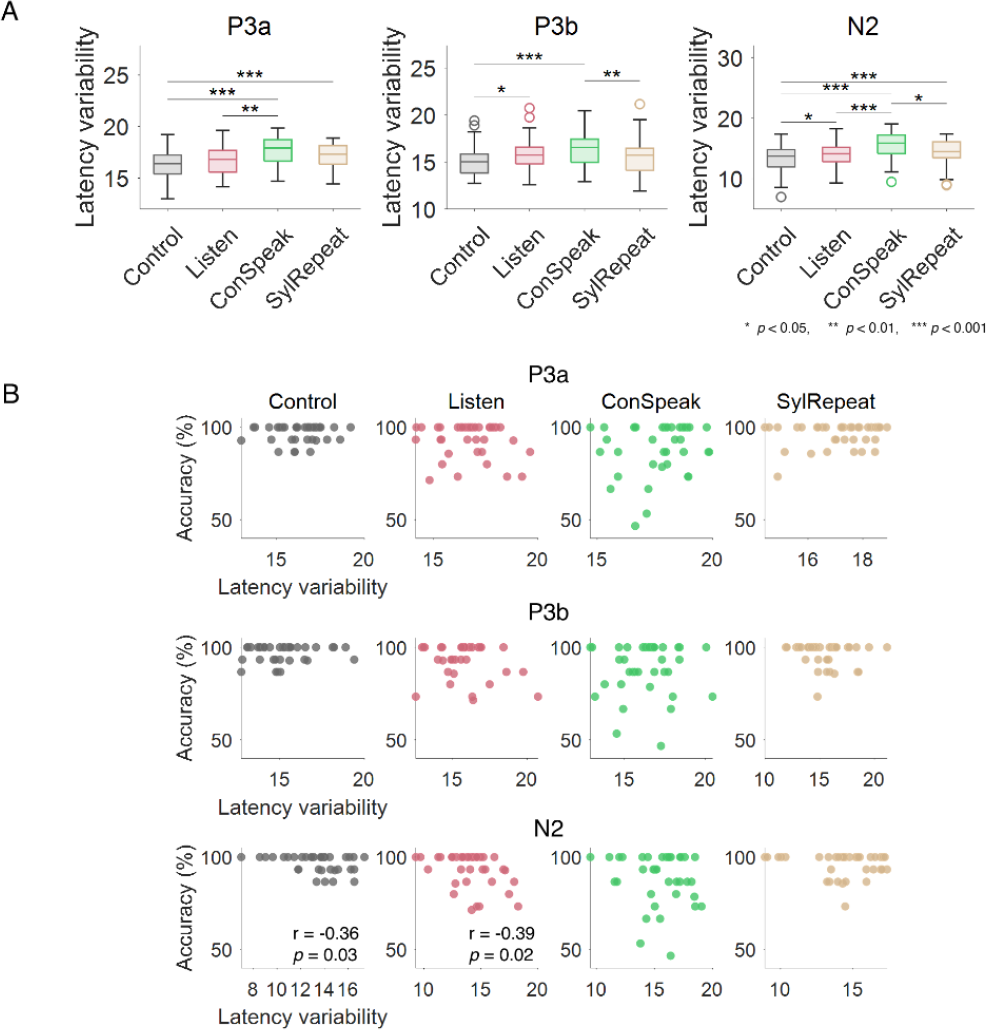
The variability of ERP latency and target classification scores. (A) The latency variability of ERP components at corresponding EEG channels. The latency variability was defined as the standard deviation of time lags at which the cross correlation of a single-trial ERP and a template ERP obtained from training data was maximized. For a detailed description, see Section 2.6.1. Statistical significance was evaluated by Friedman test with Tukey’s HSD post-hoc test, *** *p* < 0.001, ** *p* < 0.01, * *p* < 0.05. (B) The relationship between ERP latency variability and the BCI accuracies. Single dots represent individual participants. The significant correlation is depicted by the presentation of *p*-value.

We examined correlations between the latency variability and BCI accuracy across individuals for each component of P3a, P3b and N2 per condition (Figure 5(B)). For twelve combinatory cases of three components and four conditions, we found no significant correlation between the latency variability and BCI accuracy (*p* > 0.1) except for N2 in Control (*r* = -0.3581, *p* = 0.032) and Listen (*r* = -0.3914, *p* = 0.0182). This moderate negative correlation only found for N2 indicated that greater variability in latency was associated with lower BCI accuracy. Other than this, overall results collectively suggest that the BCI performance was robust to variations in latency of key ERP components.

#### 3.3.3 Standardized measurement error

The analysis of SME for the peak amplitude of ERP components revealed significant effects of conditions in most electrodes of interest, except for FC2, Cz, and Pz (Table S2). Specifically, at Fz, the SME was significantly larger in the SylRepeat condition than in Control, Listen, and ConSpeak conditions. FC1 showed a significantly larger SME in SylRepeat compared to Listen (*p* = 0.0183). At P3, the SME was significantly larger in Listen compared to Control (*p* = 0.0029). At O1, O2, and Oz, the SME was significantly larger in the ConSpeak condition than Control and Listen, with this pattern also evident in the SylRepeat condition compared to Control and Listen. In contrast, the analysis of SME for peak latency did not reveal any significant effects related to the tasks, suggesting that task variations did not substantially influence the timing variability of ERP components (Table S3).

### 3.4 Alpha band power

The alpha power modulation level, which measured discrepancy of alpha power in dual task conditions compared to the Control condition, was compared among the three conditions of Listen, ConSpeak and SylRepeat (Figure 6(A)). The alpha power modulation level was significantly lower than zero under Listen (*z* = -4.3518, *p* < 0.0001, Wilcoxon signed rank test) and SylRepeat (*z* = -3.4406, *p* < 0.0001) but not under ConSpeak (*z* = -1.0526, *p* = 0.2925). Friedman test showed a significant effect of condition on the alpha power modulation level (*χ*^2^(2) = 11.55, *p* = 0.0031). The alpha power modulation level was higher in ConSpeak than in Listen (*p* = 0.0543) and SylRepeat (*p* = 0.0029). In addition, we examined correlations between the BCI accuracy difference relative to Control (i.e., BCI accuracy in the dual task – BCI accuracy in Control) and the alpha power modulation level across individuals for each condition. A significant negative correlation appeared between the accuracy difference and the alpha power modulation level in the ConSpeak condition (*r* = -0.3663, *p* = 0.0280), but not in Listen (*r* = -0.0741, *p* = 0.6675) and SylRepeat (*r* = -0.1071, *p* = 0.5342) (Figure 6(B)).

**Figure 6.**
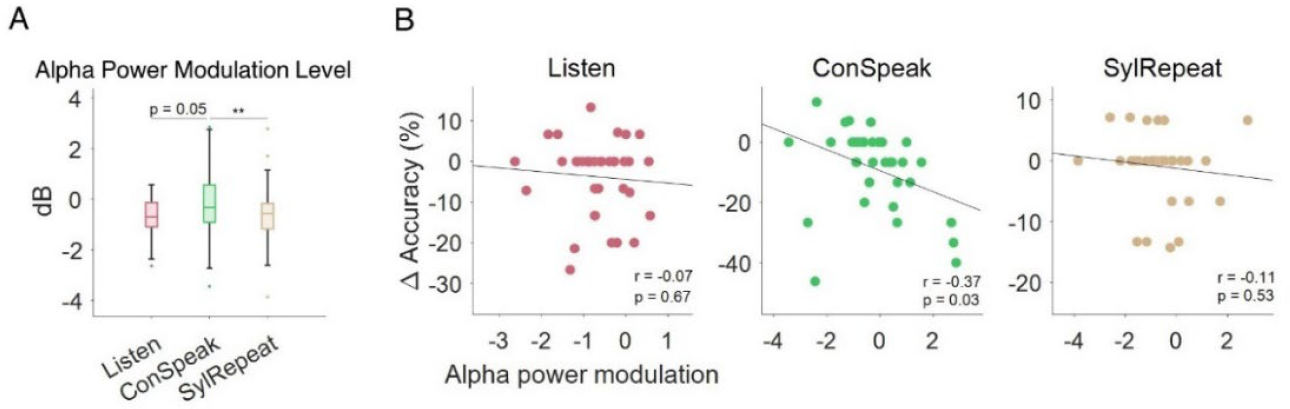
Occipital alpha power modulation level under different conditions. (A) The alpha power modulation level at occipital area, defined as the alpha power in dual task periods of each condition relative to that in Control. (B) The correlation between alpha power modulation level and the BCI accuracy difference (Accuracy_condition_ – Accuracy_Control_). A single dot indicates single participant in a specific condition. The black line is a regression line estimated using all dots in the plot.

### 3.5 Separability of classification output

The analysis of the separability of classification output, as indicated by the *d*’ values calculated for each participant (Figure 7), revealed that there were significant differences in separability between the conditions (Friedman test, *χ*^2^(3) = 48.43, *p* < 0.0001). The post hoc test showed that *d*’ was significantly smaller in ConSpeak than in all other conditions (*p* < 0.0001 compared to Control and Listen, *p* = 0.0004 compared to SylRepeat). *d’* in SylRepeat was significantly smaller than that in Control (*p* = 0.0276).

**Figure 7.**
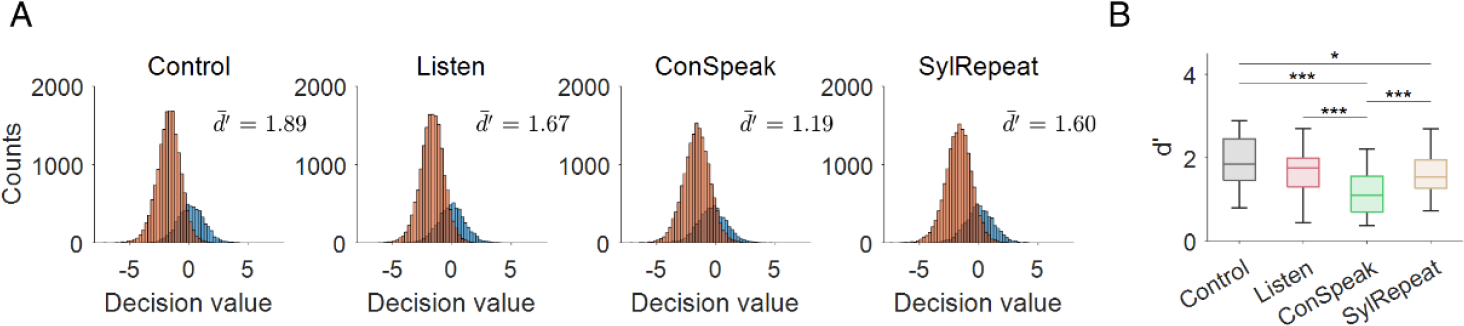
(A) Histograms representing the distribution of classification outputs under different conditions. Classification outputs are represented by the decision value that is the distance from the decision boundary of SVM to data points. 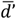 indicates averaged *d’*, a metric for separability between classes, across participants. The orange color represents nontarget trials and blue color shows target trials. (B) Comparison of *d’* among the four conditions across all 36 participants. *** *p* < 0.001, * *p* < 0.05.

## 4. Discussion

This study sought to explore the effect of conversation performed in everyday life, such as speaking or listening, on using visual ERP-BCIs. The experiment was designed to emulate real-world scenarios where users are engaged in conversation while using a BCI system. In the experiment, participants controlled ERP-BCIs to operate a smart light while speaking their opinion (ConSpeak), listening to short texts (Listen), pronouncing simple syllables as instructed (SylRepeat), or performing no other tasks (Control). The effects of these tasks were assessed based on BCI performance, the ERP components, and occipitoparietal alpha power. The results showed that ConSpeak impaired BCI accuracy the most, while Listen and SylRepeat impaired accuracy to a lesser extent than ConSpeak and a similar extent to each other. Regarding ERP components, ConSpeak reduced the amplitude of P3a, P3b, and N2 the most compared to other tasks. Among these components, N2 amplitudes showed a correlation with BCI accuracy. The level of occipital alpha power modulation was significantly greater in ConSpeak than in Listen and SylRepeat, indicating attenuated visual attention while concurrently conducting speaking task and controlling ERP-BCIs. Also, participants who exhibited higher occipital alpha power modulation levels tended to control ERP-BCIs inaccurately during speaking.

The reduced amplitudes of P3a, P3b, and N2 during speaking suggest that continuous speaking may disrupt cognitive processes essential for maintaining BCI performance, as these components are linked to attentional shift (P3a), and event categorization (P3b) [8,9,22]. In listening tasks, P3a and P3b amplitudes also decreased, though not significantly, potentially indicating less impact on BCI performance compared to speaking tasks. N2 amplitudes decreased in all dual-task conditions, most notably during speaking, highlighting N2’s close association with perceptual processing and its heightened sensitivity compared to P3a and P3b. Moreover, the correlation of N2 amplitude with BCI accuracy suggests the significant role of N2 in BCI performance, emphasizing the critical influence of visual attention level, as evidenced by N2’s linkage to visual attention [22, 23]. Hence, we speculate that distraction by conversation, especially speaking, might more likely influence visual attention than higher-order processes such as attentional shift and event categorization during the oddball task.

The occipital alpha power, another metric reflecting visual attention, further supports the effect of distraction on visual attention. In terms of the alpha power modulation level relative to the Control condition, the extent of alpha suppression appeared to be relatively smaller in the ConSpeak condition than that in other conditions (Figure 6(A)). As the extent of alpha suppression during dual tasks represents an increased level of visual attention, this result indicates a reduced level of visual attention in ConSpeak. In addition, the degree of alpha modulation exhibited a negative correlation with BCI accuracy in ConSpeak, indicating that a decreased visual attention level reflected on the increase in alpha power, may be related to decreased accuracy.

On the contrary, the occipital alpha power was suppressed during the dual task in Listen and SylRepeat relative to Control. This suppressed alpha power may indicate that visual attention needed to be elevated in the Listen and SylRepeat conditions due to the concurrent tasks. However, the N2 amplitude, which also reflects visual attention, was decreased in these conditions compared to Control. This inconsistency between the alpha power and N2 amplitudes might be partially linked to other findings that alpha oscillations do not always align with behavioural outcomes or ERPs [58] as alpha oscillations may represent cortical excitability rather than visual perception while N2 amplitudes are more directly associated with visual perception [59]. However, as overall BCI performances were relatively high in Listen and SylRepeat and not many participants showed lower accuracy in Listen and SylRepeat than in Control, it remains elusive to interpret EEG data in relation to BCI performance from the results of Listen and SylRepeat.

Our investigation into latency variability of ERP components highlights its impact on BCI performance, marking trial-by-trial variability as a cognitive function indicator. Conditions like brain trauma and ADHD show increased latency variability, often correlating negatively with BCI performance [60, 61]. In our study, ConSpeak notably increased latency variability across all ERP components, suggesting significant neural processing inconsistency across trials. Lesser but noticeable increases were seen in Listen and SylRepeat tasks. A moderate negative correlation between N2 latency variability and BCI accuracy was observed in the Listen task, suggesting auditory attention’s role in BCI performance. Yet, no significant correlations were found in ConSpeak and SylRepeat, indicating distractions may increase latency variability without a direct link to BCI performance.

While we found that conversation tasks influenced primary ERP components in the oddball paradigm, it is noteworthy that the peak amplitude and latency variability of ERP components were not directly utilized for classification in our ERP-BCIs. Therefore, after feature extraction, the classification outputs were analyzed for class separability to determine the impact of task conditions on the classification process. Similar to BCI accuracy, the separability of classification outputs was found to be the lowest in ConSpeak, while Listen showed no significant difference from Control. However, unlike BCI accuracy, the separability decreased significantly in SylRepeat compared to Control. This observation confirms that the distraction effect of speaking tasks, identified in the ERP analysis, was also reflected on classification. Although the lower separability in SylRepeat could be attributed to artifacts, the absence of a significant difference in BCI accuracy between them suggests that the impact of artifacts might not be large enough to affect BCI performance adversely.

One unexpected result is that P3b was reduced in both ConSpeak and SylRepeat compared to Control. This conflicts with the BCI performance results where the performance of BCI was higher in SylRepeat than in ConSpeak. We speculate that the decreased P3b amplitude might not only reflect attentional processes but also reflect other cognitive processes related to speech production. Speech production is a complex task that involves a number of cognitive, linguistic, and sensorimotor processes jointly working in an integrated manner [62]. Also, the parietal area, where P3b typically appears, is involved in sensorimotor integration [63] as well as the intention to speak [64]. As such, the sensorimotor integration and the intention to speak during speech production might have generated overlapped neural activity in parietal areas, affecting the P3b amplitude. However, further studies are required to investigate the potential effect of speech production on the P3b amplitude.

As ConSpeak and SylRepeat both involves facial movement, we evaluated the possible impact of artifacts on ERPs. The analysis of SME on ERPs demonstrated the impact of artifacts on ERPs in ConSpeak and SylRepeat, with no difference between ConSpeak and SylRepeat. This result suggests that the artifacts from facial movements might lower the quality of ERPs. However, ERP peak amplitudes were different between ConSpeak and SylRepeat especially at occipital areas despite similar movement-related artifacts. Also, the ERP amplitudes at diverse electrodes were not different between SylRepeat and Listen (Figure 4(A)), where no facial movement was involved in Listen. Similarly, no difference in latency variability was found between Listen and SylRepeat while latency variability was different between ConSpeak and SylRepeat at parietal (P3b) and occipital areas (N2) (Figure 5(A)). Furthermore, both data separability and BCI accuracy were different between ConSpeak and SylRepeat, but not different between Listen and SylRepeat (Figures 2(A) and 7(B)). All these observations indicate that decreased ERP quality due to facial movements may not be a main factor for a decrease in BCI performance in ConSpeak.

Subjective assessments using NASA TLX and RSME did not align with the BCI performance results. The mental effort reported by participants was the highest during ConSpeak, followed by Listen and SylRepeat, with the control task requiring the least effort. Also, the difference between ConSpeak and Listen was not significant, while accuracy in ConSpeak was significantly lower than that in Listen. This indicates that the subjective assessment of the degree of mental efforts might not be directly reflected in the BCI performance.

We observed that the ConSpeak condition yielded the lowest BCI performance, accompanied by the lowest ERP amplitudes and highest alpha power modulation. The reduced ERP amplitudes and increased alpha power during speaking could be indicative of a diversion of visual attention away from the BCI task, thereby decreasing the BCI performance. In contrast, BCI performance as well as ERP amplitudes in Listen were not significantly decreased compared to those in Control while participants reported a higher mental effort level in Listen than in Control. These results are in line with [32], which reported that attending to background speech required a higher cognitive workload, but the auditory BCI performance and the P3 amplitude did not significantly degrade. We speculate that listening to and comprehending auditory contents people encounter in daily life is not distracting enough to cause a significant reduction in the ERP amplitudes and BCI performance.

While this study provides valuable insights into the distraction effect of dual tasks on ERP-BCIs, there exist several limitations to be addressed in the future. This study focused solely on healthy undergraduate students, leaving the generalizability of the findings over different populations questionable. Additionally, even though the speaking and listening tasks were designed to mimic the real-world scenario, they lacked some factors of real-world conversation such as turn-taking, which is crucial in conversations and requires speech planning activities while listening to an interlocutor speaking [65–67]. Lastly, we only used BCIs using visual stimuli and have identified the possible influence on visual attention. As such, it is unclear whether the findings of this study can be applicable to BCIs with other sensory modalities. Future research may explore the distraction effects in different populations and with different stimulus types to validate our findings using more complex conversation tasks.

In conclusion, our study’s findings indicate that concurrent engagement in speaking tasks can affect ERP-BCI performance. This aligns with previous research on the impact of concurrent tasks and multitasking on ERP-BCIs [24–27]. Language processing, especially speaking, which requires substantial cognitive resources, seems to interfere with the attentional processes necessary for BCI control. An encouraging finding is that despite a decrement in the BCI accuracy attributed to the distractive effect of the speaking task, the mean accuracy remained above 80%. This suggests the viability of employing the BCI system in real-life environments subject to diverse distractions. This study may contribute to understanding the real-world challenges of using ERP-BCIs with daily activities. The results also highlight the need to consider the associated attentional mechanisms when designing BCIs for practical applications. Further research could delve into developing adaptive BCI systems that can dynamically adjust to users’ attentional states and tasks, ensuring more robust use of BCIs even in attention-demanding circumstances.

## Acknowledgements

This research was supported by the Challengeable Future Defense Technology Research and Development Program through the Agency For Defense Development (ADD) funded by the Defense Acquisition Program Administration (DAPA) in 2022(No.915061201).

