## Supplementary material for "Distraction Impact of Concurrent Conversation on Event-Related Potential Based Brain-Computer Interfaces": Table S1

**Table S1**. Mean and Friedman test results of ERP peak latency

| **Channel** | **Peak Latency (ms)** | | | | **Chi-square** | ***p*-value (corrected)** |
| --- | --- | --- | --- | --- | --- | --- |
|  | **Control** | **Listen** | **ConSpeak** | **SylRepeat** |  |  |
| Fz | 227.44 | 221.72 | 198.61 | 224.17 | 12.2727 | 0.0650 |
| FC1 | 229.11 | 224.22 | 211.50 | 221.44 | 4.1102 | 0.6579 |
| FC2 | 220.61 | 221.56 | 215.56 | 225.44 | 0.7328 | 0.8655 |
| Cz | 223.28 | 219.06 | 209.33 | 220.61 | 1.8102 | 0.8218 |
| P3 | 345.22 | 359.44 | 354.72 | 360.44 | 2.3898 | 0.8218 |
| Pz | 334.17 | 348.11 | 342.34 | 341.94 | 5.6071 | 0.6579 |
| P4 | 325.72 | 342.78 | 335.17 | 334.94 | 2.1404 | 0.8218 |
| O1 | 214.65 | 213.76 | 204.00 | 210.00 | 1.0599 | 0.8655 |
| Oz | 211.33 | 207.83 | 201.06 | 212.83 | 3.9845 | 0.6579 |
| O2 | 212.06 | 207.43 | 194.86 | 211.89 | 1.6084 | 0.8218 |
