## Supplementary material for "Distraction Impact of Concurrent Conversation on Event-Related Potential Based Brain-Computer Interfaces": Table S2

**Table S2**. Mean and Friedman test results of standardized measurement error (SME) of peak amplitude

| **Channel** | **SME of Peak Amplitude** | | | | **Chi-square** | ***p*-value (corrected)** |
| --- | --- | --- | --- | --- | --- | --- |
|  | **Control** | **Listen** | **ConSpeak** | **SylRepeat** |  |  |
| Fz | 0.8518 | 0.8417 | 0.8400 | 0.8940 | 14.3667 | 0.0061 |
| FC1 | 0.7430 | 0.7204 | 0.7648 | 0.7801 | 9.4333 | 0.0344 |
| FC2 | 0.7446 | 0.7319 | 0.7752 | 0.7835 | 7.2000 | 0.0731 |
| Cz | 0.7154 | 0.6851 | 0.7104 | 0.7363 | 8.2333 | 0.0518 |
| P3 | 0.7443 | 0.7047 | 0.7352 | 0.7388 | 12.6000 | 0.0112 |
| Pz | 0.7583 | 0.7293 | 0.7326 | 0.7569 | 4.4400 | 0.2177 |
| P4 | 0.7568 | 0.7582 | 0.8123 | 0.8065 | 10.6333 | 0.0231 |
| O1 | 0.9734 | 0.9518 | 1.1471 | 1.1012 | 45.0000 | 4.6263$\times$10^-9^ |
| Oz | 0.9440 | 0.9313 | 1.0636 | 1.0684 | 35.3000 | 3.5093$\times$10^-7^ |
| O2 | 0.9637 | 0.9556 | 1.1292 | 1.1114 | 45.4457 | 4.6263$\times$10^-9^ |
