## Supplementary material for "Distraction Impact of Concurrent Conversation on Event-Related Potential Based Brain-Computer Interfaces": Table S3

**Table S3**. Mean and Friedman test results of standardized measurement error (SME) of peak latency

| **Channel** | **SME of Peak Latency** | | | | **Chi-square** | ***p*-value (corrected)** |
| --- | --- | --- | --- | --- | --- | --- |
|  | **Control** | **Listen** | **ConSpeak** | **SylRepeat** |  |  |
| Fz | 27.6435 | 27.6486 | 27.7035 | 27.6986 | 0.1667 | 0.9828 |
| FC1 | 27.6855 | 27.6241 | 27.8169 | 27.6759 | 5.1333 | 0.8115 |
| FC2 | 27.6986 | 27.7215 | 27.6473 | 27.8583 | 5.8333 | 0.8115 |
| Cz | 27.7846 | 27.7317 | 27.6563 | 27.6932 | 2.5333 | 0.8124 |
| P3 | 28.7515 | 28.5398 | 28.8298 | 28.9306 | 0.5000 | 0.9828 |
| Pz | 28.8222 | 28.8411 | 28.6782 | 28.7072 | 1.1829 | 0.9828 |
| P4 | 28.7256 | 28.6555 | 28.7328 | 28.7608 | 0.2333 | 0.9828 |
| O1 | 27.5909 | 27.6450 | 27.6422 | 27.6528 | 3.1059 | 0.8124 |
| Oz | 27.6736 | 27.7585 | 27.6676 | 27.6082 | 2.4333 | 0.8124 |
| O2 | 27.7896 | 27.7588 | 27.6826 | 27.8132 | 2.5543 | 0.8124 |
